# Transcranial Static Magnetic Stimulation Dissociates the Causal Roles of the Parietal Cortex in Spatial and Temporal Processing

**DOI:** 10.1101/2025.11.16.688651

**Authors:** Masakazu Sugimoto, Ikko Kimura, Masamichi J. Hayashi

**Author notes:** Corresponding author: Center for Information and Neural Networks (CiNet), Advanced ICT Research Institute, National Institute of Information and Communications Technology, 1-4 Yamadaoka, Suita, 565-0871, Japan, (Masamichi J. Hayashi).

## Abstract

Accurate time estimation is essential for optimizing our perception and actions. Previous neuroimaging and transcranial magnetic stimulation (TMS) studies have suggested that the right inferior parietal lobule (IPL) and supplementary motor area (SMA) are involved in time perception. However, it remains inconclusive whether the activity in these regions is crucial for time perception, partly due to the possible spread of TMS effects across anatomically connected brain regions. Such a remote effect is less likely to happen with transcranial static magnetic stimulation (tSMS), as the static magnetic field is expected to modulate the firing threshold of neurons rather than directly triggering an action potential. In this study, we aimed to determine the causal relevance of local activities in the right IPL and the SMA for temporal processing using tSMS. 48 human volunteers (26 males and 22 females) participated in the study. We measured duration discrimination thresholds, along with orientation discrimination thresholds, using staircase methods before and during the administration of tSMS over the IPL/SMA. Our results indicated no significant changes in duration discrimination thresholds in either the IPL or SMA conditions. In contrast, we observed an improvement in orientation discrimination thresholds in the IPL condition. This improvement correlated with individual differences in the distance between the scalp and the IPL. Overall, our findings demonstrate a causal involvement of the IPL in orientation processing. The correlation between the effects of tSMS and the scalp-to-target distance suggests that the efficacy of tSMS may be sensitive to the magnetic field strength.

**Significance Statement:** Accurate time estimation is essential for optimizing our perception and actions. While prior studies using transcranial magnetic stimulation (TMS) investigated a role of inferior parietal lobule (IPL) and supplementary motor area in spatiotemporal processing, the results have been inconclusive because TMS can affect anatomically connected areas. Here, using transcranial static magnetic stimulation (tSMS), we examined the causal relevance of local activities in these areas. Our findings showed that tSMS over the IPL did not alter duration discrimination but, unexpectedly, significantly enhanced orientation discrimination, with the degree of improvement correlating with individual anatomical differences. These results provide new insights into the neural basis of spatial and temporal processing and emphasize the potential of tSMS as a localized neuromodulation technique of interest.

## Introduction

Precise estimation of time is essential for adapting to an ever-changing environment and optimizing our actions. For instance, in daily activities such as conversation and dancing to a beat, the ability to accurately estimate and predict the timing of external events enables smooth and effective interactions (Teki et al., 2011; Magyari et al., 2014, 2017; Ross et al., 2016).

Recent functional magnetic resonance imaging (fMRI) studies have shown that multiple brain regions are involved in temporal processing in the range of hundreds of milliseconds to about one second (Wiener et al., 2010b; Coull and Droit-Volet, 2018; Naghibi et al., 2024). A meta-analysis of neuroimaging studies reported that perceptual, subsecond timing tasks engage a distributed network of brain regions, including the supplementary motor area (SMA), inferior parietal lobule (IPL), inferior frontal gyrus (IFG), and basal ganglia (Wiener et al., 2010b). Among these areas, the right IPL and SMA have been highlighted as they seem to contain neural populations tuned for specific durations (Hayashi et al., 2015; Protopapa et al., 2019; Harvey et al., 2020). More recent research has associated activity in the right IPL with individual differences in time discrimination performance (Hayashi et al., 2018) and perceived duration (Hayashi and Ivry, 2020), suggesting its relevance to our subjective experience of time.

Consistent with these neuroimaging findings, several studies have reported that transcranial magnetic stimulation (TMS) applied over the right IPL impairs time discrimination (Bueti et al., 2008; Riemer et al., 2016) or biases perceived durations (Wiener et al., 2010a, 2012). However, it remains unclear whether the modulation of local activity in the area directly affects timing performances or operates through a remote effect, where TMS influences neuronal activity in anatomically connected regions (i.e., remote effect) (Rafique et al., 2015; Bergmann et al., 2021; Wang et al., 2024). This is of particular concern given that a recent functional connectivity study reported that the parietal cortex drives duration-selective activities in several brain regions, including the SMA, visual cortex, IFG, and cerebellum (Protopapa et al., 2023). In line with this finding, a TMS-EEG study reported that event-related potentials in frontocentral electrodes around the SMA were modulated by TMS applied over the right IPL and correlated with changes in timing performance (Wiener et al., 2012). These studies suggest that, even if TMS produces measurable behavioral effects, it may be premature to attribute these effects to changes in local activity beneath the TMS coil.

A potentially useful neuromodulation technique for determining the causal role of specific brain regions is transcranial static magnetic stimulation (tSMS) (Oliviero et al., 2011). tSMS is a non-invasive brain stimulation method that modulates the firing threshold of neurons in the cortex by placing a neodymium magnet on the scalp (Sinha et al., 2024). Although the precise mechanism is not fully understood, it is believed that a static magnetic field of approximately 120–200 mT affects sodium and chloride ion channels, possibly by deforming the lipids in the cell membrane (Rosen, 2003; Hernando et al., 2020; Sinha et al., 2024). These alternations in ion influx may lead to inhibitory effects on neural activity within the area exposed to the static magnetic field. Since tSMS does not trigger action potential of neurons, it may offer advantages over TMS in producing effects that are relatively localized and with minimal remote influences on anatomically connected brain regions.

To determine whether the local activities of IPL are crucial for temporal processing, here, we examined the influence of tSMS over the region on the duration discrimination task. If the area contains duration-tuned neuronal populations that are crucial for subsecond timing, we predicted that suppressing the IPL through tSMS should alter duration discrimination performance. In addition to targeting the IPL, we included stimulation over the SMA as an exploratory feasibility test for the application of tSMS. This decision was motivated by prior timing studies reporting limited effects of TMS over the SMA (Del Olmo et al., 2007; F. Giovannelli et al., 2014; Méndez et al., 2017), potentially because timing-related responses are localized to deeper, medial portions of the SMA, as demonstrated by previous fMRI studies (Hayashi et al., 2018; Protopapa et al., 2019). Participants also performed an orientation discrimination task. For this task, we predicted that tSMS over neither the IPL nor the SMA would alter performance, because a previous neuroimaging study showed that while duration information was decoded from the IPL and, albeit less clearly, from the SMA, orientation information was not evident in these regions (Hayashi et al., 2018).

## Materials and Methods

### Participants

A total of 48 healthy, right-handed adult volunteers participated in our experiments (22 females and 26 males; age = 22.1 (mean) ± 1.4 (SD) years). The sample size of 48 participants was determined a priori based on statistical power consideration. Specifically, detecting a moderate correlation with adequate statistical power requires a minimum of 46 participants (Bujang and Baharum, 2016). Given that our experimental design required the total number of participants to be a multiple of four to allow for counterbalancing, we set the final sample size to 48. This sample size is comparable to, and in some cases larger than, those used in previous tSMS studies targeting the SMA (Pineda-Pardo et al., 2019; Guida et al., 2023).

Participants were recruited through the institutional online recruitment system from July 15, 2021, to April 14, 2023. All participants had normal or corrected-to-normal vision and no history of neurological and psychiatric disorders. This study was conducted in accordance with the Declaration of Helsinki. Participants received detailed written explanations of the experimental procedures and safety measures, and they provided informed consent. The protocol was approved by the institutional ethics and safety committees.

### Task and stimuli

Visual stimuli were generated using MATLAB (MathWorks, Natick, MA, USA) and presented on a gamma-corrected CRT monitor (Multiscan G520; Sony Inc., Tokyo, Japan) with a refresh rate of 100 Hz. In each trial, a standard stimulus and a comparison stimulus were sequentially presented at the center of the screen, with an inter-stimulus interval (ISI) of 1.0 s (Fig. 1). Following the two visual stimuli, a red fixation point was presented to indicate the response period. The red fixation point remained on the screen until the participant responded. The subsequent trial began immediately after the response. Participants were instructed to maintain their gaze on the fixation point throughout the experiments.

**Fig. 1.**
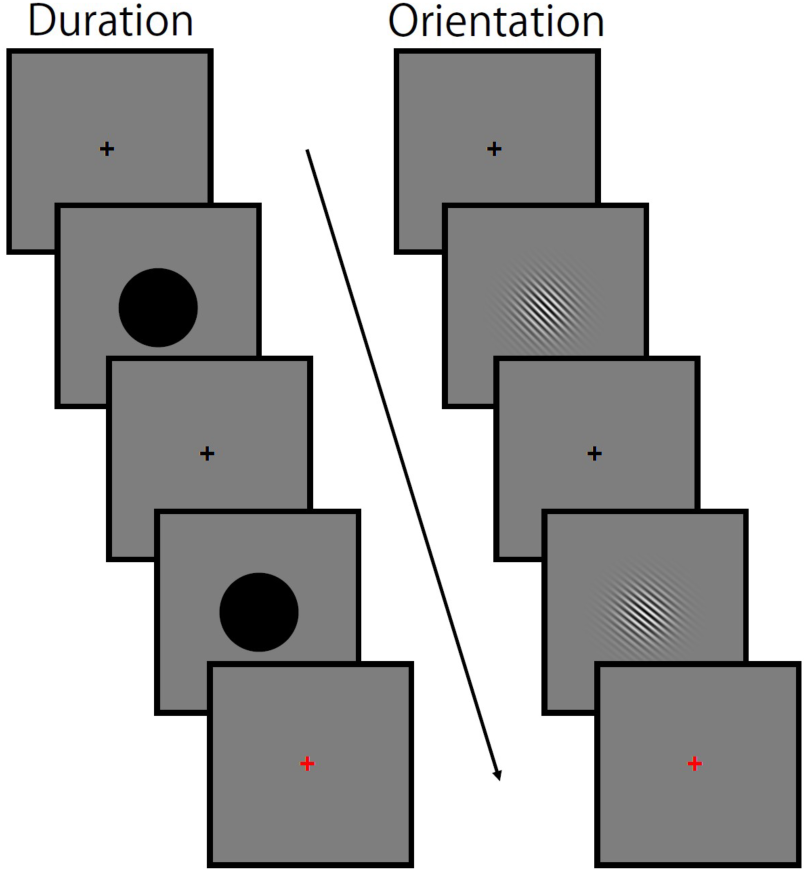
Stimulus sequence. In the duration task (left), two visual stimuli (black disks) were presented sequentially. Participants judged whether the duration of the second disk was shorter or longer than that of the first. In the orientation task (right), two Gabor patches were presented sequentially. Participants judged whether the orientation of the second stimulus was rotated clockwise or counterclockwise compared to the first. A response was required in every trial during the response period, indicated by a red fixation point.

In the duration task, the standard and comparison stimuli were black disks (diameter 5.4 degrees), and participants judged whether the second stimulus was shorter or longer than the first. In the orientation task, the standard and comparison stimuli were sinusoidal Gabor patches (100% contrast, spatial frequency of 1.6 cycles/degree, Gaussian envelope SD of 2.0 degrees, diameter ∼12 degrees). Participants judged whether the second stimulus was rotated clockwise or counterclockwise relative to the first.

A one-up-three-down staircase procedure was employed to measure participants’ thresholds in both the duration and orientation discrimination tasks. In the duration discrimination task, the duration of the standard stimulus was set to 0.5 s, while the duration of the comparison stimulus was 0.5 s + T (initial T = 0.4 s). In the orientation discrimination task, the standard orientation of the sinusoidal Gabor patches was 45° from vertical while that for the comparison was 45° + T (initial T = 12°). The stimulus duration for each Gabor patch was 0.06 s. The order of presentation for the standard and comparison stimuli was randomized across trials.

Until the first three reversals, T was multiplied by 1.5 (increased) when the response was incorrect and divided by 1.5 (decreased) when the response was correct for both tasks. After obtaining three reversal points, T was multiplied by 1.3 when the response was incorrect or divided by 1.3 when the response was correct for three consecutive trials. To mitigate any potential confounds related to experiment duration (e.g., fatigue), the measurement durations were fixed at 270 s for the duration task and 215 s for the orientation task based on pilot experiments. Participants responded by pressing either the “J” key (for shorter or counterclockwise) or the “K” key (for longer or clockwise) with their right index and middle fingers, respectively. Participants were explicitly instructed to respond as accurately as possible and not to count time during the duration task.

### Target localization for tSMS

We employed a tSMS, a cylindrical neodymium magnet, with a diameter of 60 mm and a thickness of 30 mm (MAG 60 r+, Neurek SL., Toledo, Spain). The magnetic field strength was 0.511 T at the surface of the magnet. The right IPL (MNI coordinates x, y, z = 66, −34, 34) and SMA (x, y, z = 2, 0, 70) were selected as the stimulation sites based on preceding fMRI studies (Hayashi et al., 2018; Hayashi and Ivry, 2020). To determine the individual’s positions of the stimulation sites, we collected T1-weighted images for each participant with a magnetization-prepared rapid acquisition with a gradient echo (MPRAGE) sequence using a 3T MRI scanner (Siemens Inc., Erlangen, Germany), MAGNETOM Prisma (0.9 mm isotropic resolution, TR = 2700 ms, and TE = 3.68 ms) or MAGNETOM Prisma Fit (0.8 mm isotropic resolution, TR = 2500 ms, and TE = 2.18 ms). For each participant, we localized the right IPL and SMA by converting the MNI coordinates into positions within that individual’s structural T1-weighted image. Using statistical parametric mapping (SPM12) software implemented in MATLAB, we calculated an affine transformation matrix to align the individual’s T1-weighted image with the standard brain (MNI template). Subsequently, we used the inverse of the transformation on the neuronavigation system (Brainsight, Rogue Research Inc., Montreal, Quebec, Canada) to determine the positions of the right IPL and SMA within the participant’s structural image.

### Experimental Procedure

We conducted two sessions of experiments on different days: one targeting the IPL and the other targeting the SMA. The interval between sessions ranged from 1 to 27 days. Before each session, we aligned the MR images with the participant’s head and marked the scalp position for placing the magnet using the Brainsight navigation system. Participants completed 10 practice trials for both the duration and orientation discrimination tasks.

Experiments consisted of baseline and test blocks, with each block lasting approximately 20 min. In these blocks, participants performed duration and orientation discrimination tasks two times each (Fig. 2A). Following baseline threshold measurements, which were taken without applying tSMS, we fixed the magnet over the targeted region using a custom-assembled clamp with a sliding arm (Slick Co., Ltd., Tokyo, Japan), ensuring that the center of the magnet’s bottom surface aligned with the marked position on the scalp. A chin rest was used to stabilize the participant’s head position. Based on a previous report showing suppression of cortical excitability following administration of tSMS for 30 min (Dileone et al., 2018; Takamatsu et al., 2021), we positioned tSMS over the IPL or SMA for 30 min prior to the test block and maintained the position throughout the test block (Fig. 2B).

**Fig. 2.**
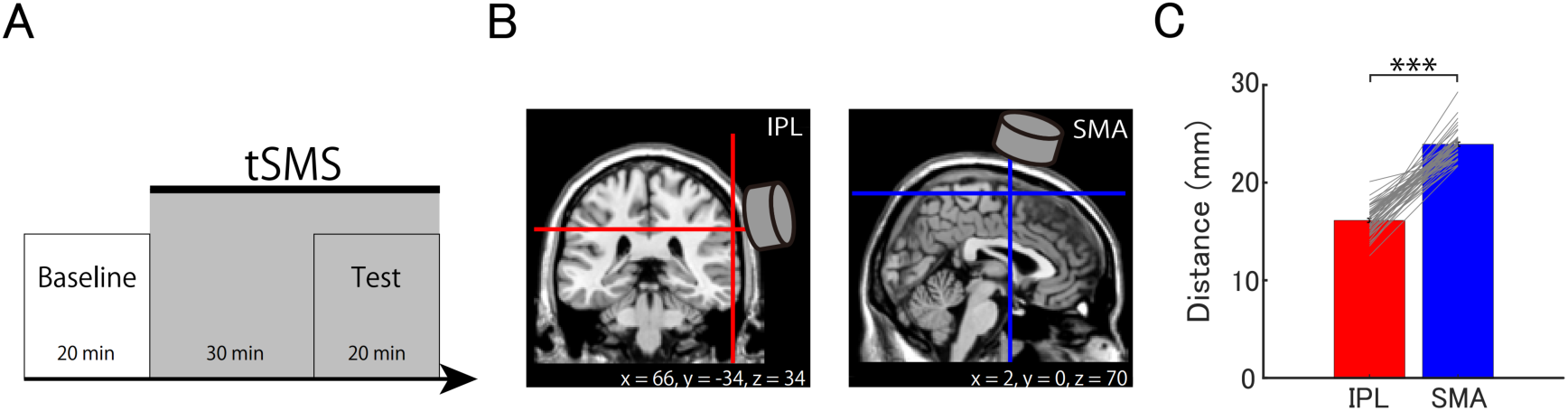
Experimental procedure and scalp-to-target distance. (A) Timeline of the experiment. During both the baseline and test blocks, participants performed duration and orientation tasks, each repeated two times. tSMS was applied for approximately 50 min, beginning 30 min before and continuing throughout the 20 min test block. (B) Target regions. We identified the target regions, right IPL and SMA, based on MNI coordinates from previous studies. The figure illustrates the target regions on coronal (left) and sagittal (right) views of a standard brain image. (C) The average distance from the scalp to the IPL and the SMA across participants. The red bar represents the mean distance to the IPL, while the blue bar represents the mean distance to the SMA. Error bars indicate the standard error. Gray lines represent individual data. *** p < 0.001

Participants were randomly assigned to one of two groups: one group received tSMS over the IPL in the first session and over the SMA in the second, while the other group was tested in the opposite order. The task order for the baseline and test blocks was counterbalanced within each group, with half of the participants performing the duration task followed by the orientation task, and the other half performing the tasks in reverse order. For each participant, the task order remained consistent across both sessions.

### Data Analysis

To determine the discrimination thresholds with the staircase procedure, we averaged the last six reversal points from the collected data (Hayashi et al., 2014). The threshold values were then averaged between two measurements to establish an individual’s discrimination threshold. If the number of reversal points in one of the two measurements was less than eight, we excluded the data and used the threshold from the other measurement as the participant’s discrimination threshold.

To assess any potential changes in the baseline threshold across sessions, we compared the average baseline threshold values between the first and second sessions using two-tailed paired t-tests (α = 0.05), with adjustments for multiple comparisons using the Bonferroni correction. Also, to evaluate the stability of individual differences in baseline discrimination thresholds, we calculated Pearson correlation coefficients between the baseline thresholds obtained in the first and second sessions. To investigate the changes in discrimination thresholds induced by tSMS, we performed two-tailed one-sample t-tests (α = 0.05) on the difference between the test and baseline thresholds, applying the Bonferroni correction for multiple comparisons across two target brain regions.

Finally, we examined whether the magnitude of the tSMS effects depends on the distance between the targeted cortical area and the magnet. We calculated the Pearson correlation coefficient between the distance from the scalp to the stimulation site (either the IPL or SMA) and the change in discrimination threshold between the test and baseline blocks. The distance was measured using the ruler tool in the Brainsight navigation system. We tested the significance of correlations (two-tailed, α = 0.05). Differences in correlation coefficients between tasks were also tested by bootstrap resampling (10,000 samples) using the Robust Correlation Toolbox implemented in MATLAB (Pernet et al., 2013).

All statistical analyses, except for the bootstrap resampling, were performed using JASP (https://jasp-stats.org/). Bayes factors (BF) were calculated for all tests to quantify the relative evidence supporting the null versus the alternative hypothesis. All Bayesian analyses were also performed in JASP.

## Results

### Reliability of the baseline threshold

Among the collected data from the 48 participants, threshold data from eleven measurements in the IPL sessions (four baseline and three test threshold measurements for the duration task, and one baseline and three test thresholds measurements for the orientation task) and six measurements in the SMA sessions (three baseline and two test thresholds measurements for the duration task and one test threshold measurements for the orientation task) were disregarded due to the insufficient number of reversal points. For these blocks, the individual’s baseline or test threshold was determined based on the threshold value obtained in the other measurement (see *Data analysis* section in the Materials and Methods).

Preceding to our main analyses, we examined whether the baseline thresholds differed between the first and second sessions. As expected, a paired t-test revealed that the baseline thresholds for duration discrimination were not statistically different (t (47) = 0.258, p = 1.000, Cohen’s d = 0.037, BF = 0.162) (Fig. 3A, left). In contrast, baseline thresholds for orientation discrimination were significantly lower in the second session compared to the first session (t (47) = 4.244, p < 0.001, Cohen’s d = 0.613, BF = 226.832) (Fig. 3B, left). This result suggests that, while baseline thresholds were maintained in the duration task, the threshold for the orientation task was improved as the experiment progressed.

**Fig. 3.**
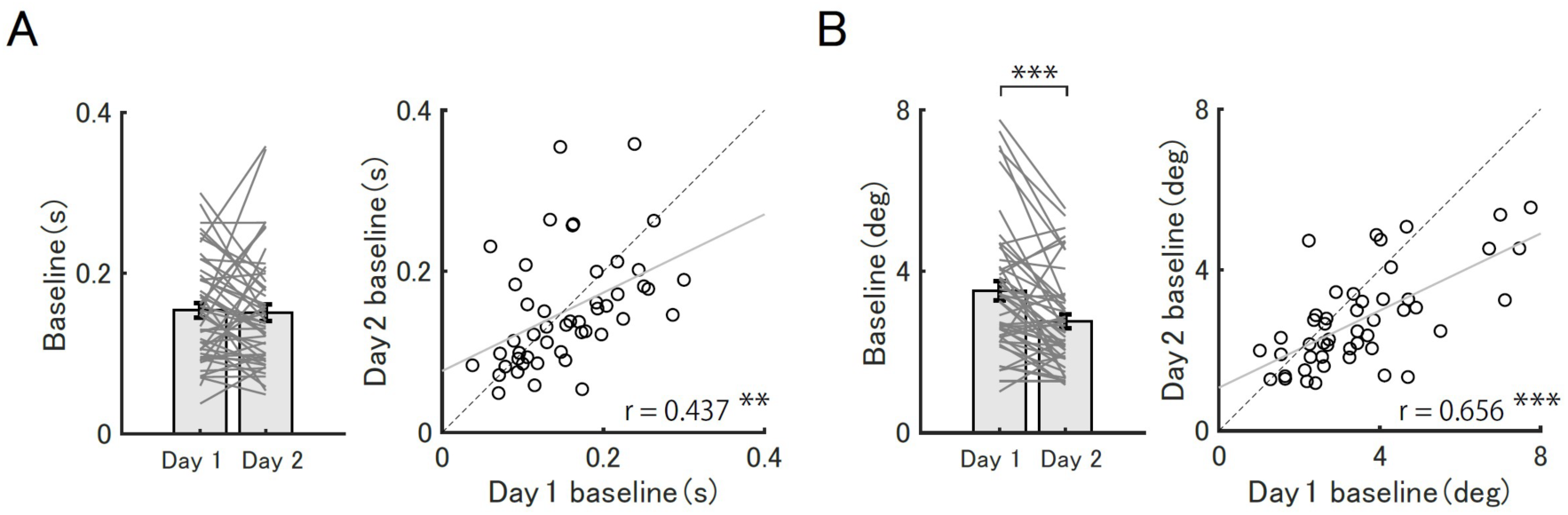
Baseline thresholds. Baseline thresholds for the first (Day 1) and second sessions (Day 2) for the duration task (A, left) and for the orientation task (B, left). Each bar represents the average discrimination thresholds, with error bars indicating the standard error. The gray lines indicate the change in individual thresholds between Day 1 and Day 2. ** p < 0.01, *** p < 0.001, corrected for multiple comparison. Correlations between thresholds for Day 1 and Day 2 for the duration task (A, right) and the orientation task (B, right). The black circles represent individual data. The gray solid line represents the regression line. The black dashed line represents the diagonal line. *** p < 0.001

We also tested whether the individual differences in baseline thresholds were maintained across sessions. Our correlation analyses revealed that individual differences between the first and the second baseline thresholds were correlated in both duration (r = 0.437, p = 0.002, BF = 19.242) (Fig. 3A, right) and orientation tasks (r = 0.656, p < 0.001, BF = 43447.499) (Fig. 3B, right). These results indicate that, although the threshold was improved in the second session in the orientation task, individual differences in the baseline thresholds were maintained across sessions in both tasks.

### Effect of tSMS on task performances

Next, we examined whether tSMS over the IPL and SMA altered discrimination thresholds in duration and orientation tasks. A one-sample t-test for the duration discrimination threshold revealed no significant difference from the baseline, both in the IPL and SMA conditions (IPL: t (47) = −0.716, p = 0.954, Cohen’s d = −0.103, BF = 0.200; SMA: t (47) = 0.381, p = 1.000, Cohen’s d = 0.055, BF = 0.168; Fig. 4A). In contrast, the orientation discrimination threshold significantly decreased when tSMS was applied over the IPL (t (47) = −2.463, p = 0.034, Cohen’s d = −0.356, BF = 2.371; Fig. 4B). No such reduction in the orientation discrimination threshold was observed in either task when tSMS was applied over the SMA (t (47) = −0.393, p = 1.000, Cohen’s d = - 0.057, BF = 0.169; Fig. 4B).

**Fig. 4.**
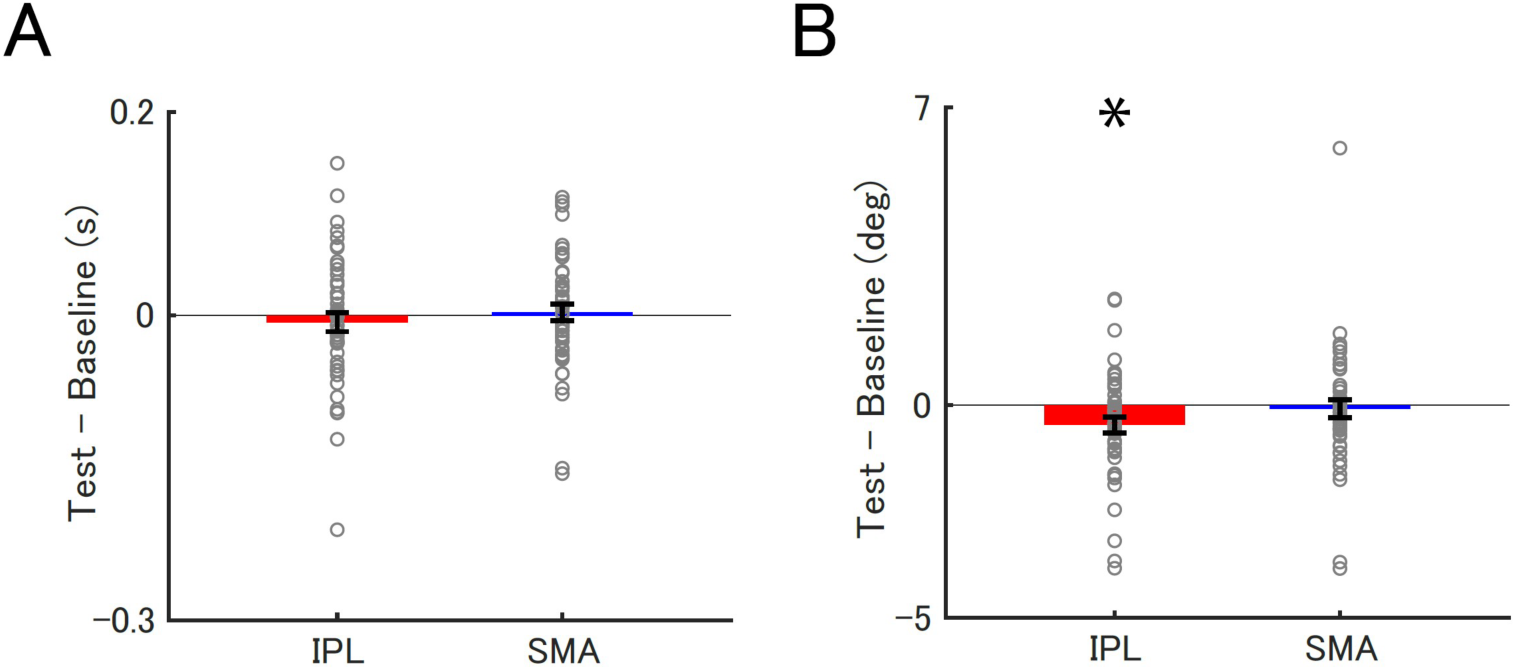
Modulation of discrimination thresholds by tSMS. Differences in the discrimination threshold between baseline and test phases (Test - Baseline) in the duration (**A**) and the orientation tasks (**B**) when tSMS was applied over the IPL (red bars) and SMA (blue bars). The gray circles indicate individual data. Each bar depicts the mean threshold difference, with error bars indicating the standard error. * p < 0.05, corrected for multiple comparisons.

### Effect of magnetic field strength on performance improvement

Since the effective magnetic field strength of tSMS depends on the distance between the magnet and target, we wondered whether the variability in the effect size of tSMS across participants is explained by the scalp-to-target distance. First, we examined whether the mean scalp-to-target distance differed between the two target regions. A paired t-test revealed that the scalp-to-target distance for the IPL (16.1 ± 0.2 mm) was significantly closer than for the SMA (24.0 ± 0.2 mm) (t (47) = −25.467, p < 0.001, Cohen’s d = −3.676, BF = 7.253 × 10^25^; Fig. 2C), indicating that magnetic field strength in the IPL was greater and probably more effective than that in the SMA.

Next, we tested whether individual differences in the scalp-to-target distance explain the effect size of tSMS. Our correlation analyses revealed that individual differences in the scalp-to-IPL distance were positively correlated with the change in orientation discrimination thresholds (r = 0.304, p = 0.036, BF = 1.526; Fig. 5B), whereas such a correlation was absent for the duration task (r = −0.029, p = 0.846, BF = 0.183; Fig. 5A). A bootstrap analysis further confirmed that the degree of correlations significantly differed between the two tasks (95% confidence interval (CI) = −0.607–-0.006, p = 0.046; Fig. 5C). For the SMA, individual differences in the scalp-to-SMA distance were not correlated with the degree of threshold modulation in either the duration (r = 0.022, p = 0.883, BF = 0.182; Fig. 5D) or orientation task (r = 0.016, p = 0.917, BF = 0.181; Fig. 5E). Our bootstrap analysis confirmed that the correlations not significantly differed between these two tasks (95% CI = −0.396–0.456, p = 0.941; Fig. 5F). These findings together suggest that the scalp-to-target distance in the IPL, which was shorter than that in the SMA, was predictive of the degree of improvement in behavioral performances, specifically for the orientation task.

**Fig. 5.**
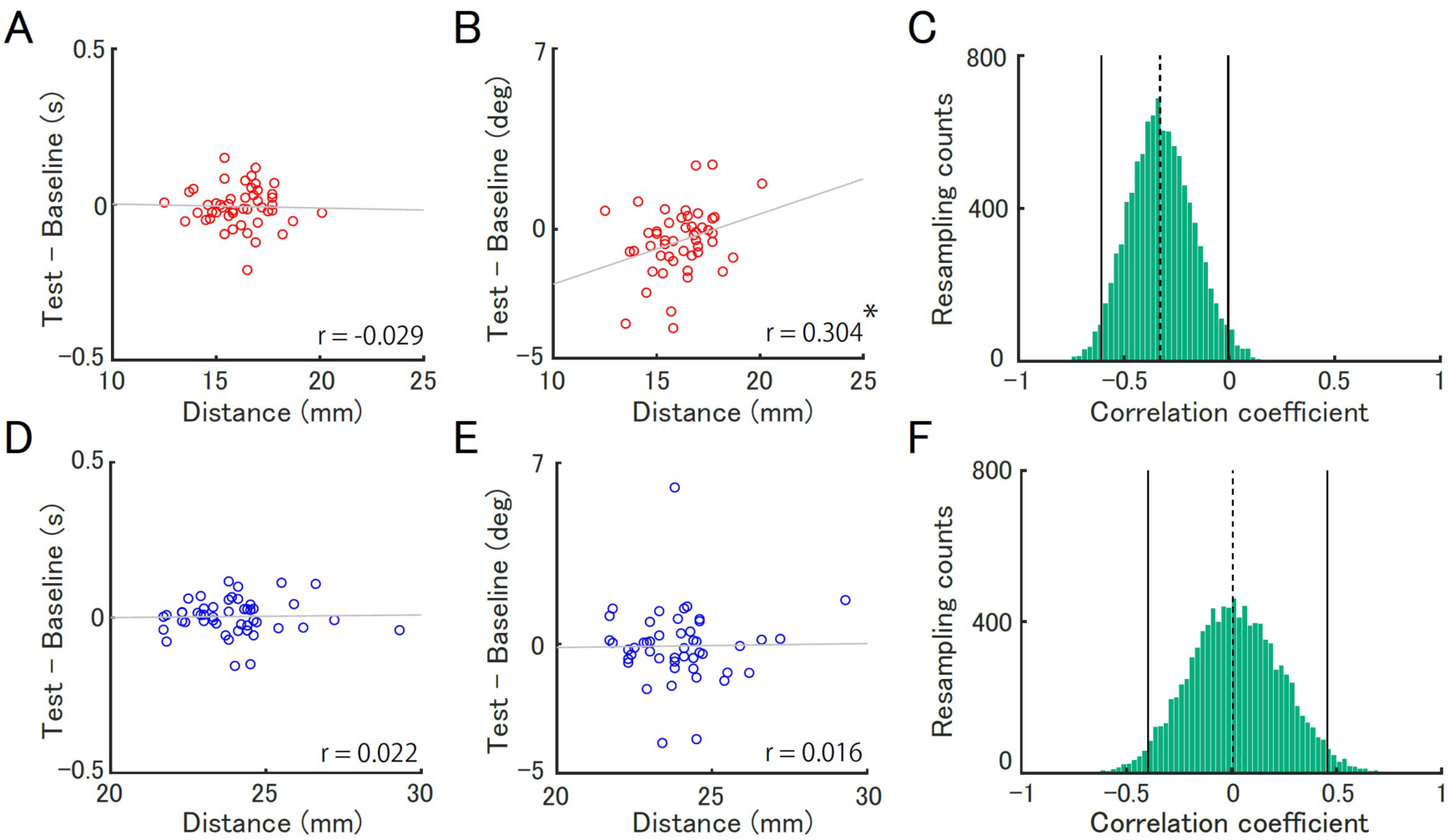
Correlation between the scalp-to-target distance and the behavioral effect of tSMS. Correlation between the scalp-to-target distance (**A and B** for IPL**, D and E** for SMA) and the change in discrimination thresholds (**A and D** for duration task, **B and E** for orientation task). Gray lines indicate regression lines and circles indicate individual data. Distribution of differences in correlation coefficients between tasks (i.e., duration - orientation task) for IPL (C) and SMA (F), estimated by the bootstrap method. The solid line represents the 95% confidence interval, and the dashed line indicates the value of differences in correlation coefficients between tasks. * p < 0.05

## Discussion

This study investigated whether the local activities in the IPL and SMA are causally involved in temporal and spatial processing by measuring duration and orientation discrimination thresholds, respectively, before and during the application of tSMS. Our within-subject experiments, which included a relatively large sample size (n = 48), showed no significant changes in duration discrimination thresholds, irrespective of the target regions. Unexpectedly, however, we found that orientation discrimination thresholds significantly improved when tSMS was applied over the IPL (but not SMA). Additionally, the degree of improvement in the orientation task correlated with individual differences in the scalp-to-target distance of the IPL.

### Lack of tSMS effect on duration task

In our experiment, performance in the duration task was not significantly altered by the tSMS applied over the IPL or SMA. This result may seem surprising given that numerous neuroimaging studies have reported the involvement of these regions in temporal processing (Coull et al., 2015; Hayashi et al., 2015, 2018; Protopapa et al., 2019; Hayashi and Ivry, 2020). The absence of a tSMS effect in our study might be related to the distributed nature of temporal representations in the brain. A lack of disorders associated with a selective impairment in timing abilities suggests that time representations might be distributed across various regions (Wiener et al., 2011). Supporting this idea, previous neuroimaging studies have consistently shown that timing information is spread across cortical areas, forming a timing network (Hayashi et al., 2018; Harvey et al., 2020; Cona et al., 2021, 2024; Protopapa et al., 2023). Therefore, we speculate that disrupting one region of this timing network may not significantly affect timing performance because the brain can flexibly utilize other reliable sources of timing information within the network to maintain behavioral performance.

Although our study found no evidence of tSMS effect on time discrimination, previous studies have reported impairments in timing performance when TMS was applied over the IPL (Bueti et al., 2008; Wiener et al., 2010a, 2012). This discrepancy may be explained by the combination of the distributed timing network idea described above and the difference in the degree of focality between tSMS and TMS methods. TMS is known to influence neuronal activity not only in the targeted areas but also in anatomically connected regions, which is referred to as the remote effect (Sack et al., 2007; Hirose et al., 2011; Watanabe et al., 2015; Siebner et al., 2022). In contrast, tSMS is believed to have a more focal effect due to its mechanism of working through the deformation of lipids in the cell membrane (Rosen, 2003; Hernando et al., 2020). Therefore, the varying degrees of focality between TMS and tSMS may account for different behavioral outcomes. Specifically, while TMS can disrupt timing performance through broader interference within the timing network, more localized disruption by tSMS might be compensated for by utilizing timing information from intact areas.

The lack of tSMS effect on the SMA could be explained by the significant distance between the SMA and the scalp. As demonstrated in the orientation discrimination task (Fig. 5B), the impact of tSMS on behavioral performance is influenced by the scalp-to-target distance. The targeted SMA was located on the medial wall of the brain, which may have rendered the magnetic field too weak to effectively disrupt cortical activity. This notion is consistent with previous TMS studies that reported no significant effect on time perception when TMS was administered over the SMA (Del Olmo et al., 2007; Fabio Giovannelli et al., 2014; Méndez et al., 2017). Our findings suggest a need for future studies to employ neuromodulation techniques that can effectively stimulate deeper areas (e.g., transcranial focused ultrasound stimulation) to better clarify the role of the SMA in duration discrimination tasks.

Although the relatively large scalp-to-target distance for the SMA suggests that the effective magnetic field reaching the cortex may have been weaker, which may potentially attenuating the impact of tSMS, previous studies have nevertheless demonstrated that tSMS applied over the SMA can modulate behavioral and physiological measures in the motor domain (Pineda-Pardo et al., 2019; Pagge et al., 2024). Importantly, these studies employed stimulation procedures broadly comparable to ours in terms of magnet size, strength, and target definition, making it unlikely that simple methodological differences alone account for the discrepancy in effectiveness across studies. Instead, the impact of tSMS on the SMA may be task dependent, reflecting differences in the neural networks recruited by the task. Specifically, prior work primarily examined motor processes such as action planning and control, in which behavioral effects may arise through modulation of interactions within the motor network, including SMA–M1 connectivity (Pineda-Pardo et al., 2019). In contrast, duration discrimination is thought to rely on widely distributed timing representations across a broader timing network (Protopapa et al., 2023). Consequently, perturbation of a single node within this network may be compensated for by intact regions, thereby reducing the likelihood that measurable behavioral effects emerge. Consistent with this interpretation, TMS studies have repeatedly reported reliable effects of SMA stimulation on motor control (Zandbelt et al., 2013; Obeso et al., 2017; Tosun et al., 2017; Berkay et al., 2018), whereas stimulation of the SMA has often yielded null effects in time perception tasks (Del Olmo et al., 2007; F. Giovannelli et al., 2014; Méndez et al., 2017). Taken together, these findings suggest that the behavioral impact of SMA stimulation may depend on the nature of the targeted task and, consequently, on the networks it engages.

### tSMS over the right IPL improved orientation discrimination performance

Orientation discrimination performance was improved when tSMS was applied over the right IPL. This finding aligns with previous studies showing that TMS over the right IPL significantly increased reaction times for detecting target orientations (Sack et al., 2007; Sack, 2009), suggesting that the IPL plays a crucial role in orientation discrimination.

The paradoxical enhancement in performance due to the inhibitory effects of tSMS can be attributed to increased lateral inhibition mechanisms of orientation-tuned neural populations in the IPL. In the visual cortex, higher GABA concentration levels have been linked to better orientation thresholds (Edden et al., 2009), likely due to the enhanced lateral inhibition in orientation-tuned neurons (Li et al., 2008; Brouwer and Heeger, 2011; Del Rosario et al., 2025). Notably, a recent fMRI adaptation study highlighted the existence of spatiotopic receptive fields for orientations in the IPL (Dunkley et al., 2016). Given that another study reported that tSMS enhances inhibitory signal mediated by GABA receptors (Nojima et al., 2015), we speculate that our tSMS application might have increased GABA concentration or sensitivity of GABA receptors associated with orientation channels in the IPL. Consequently, this rise in GABA levels could have enhanced orientation selectivity, leading to improved orientation discrimination performance.

Additionally, we found that participants with shorter scalp-to-target distances showed greater improvements in their orientation discrimination performance. This finding suggests that scalp-to-target distance could be a useful metric for evaluating the effectiveness of tSMS. For other neuromodulation techniques, such as TMS, simulating induced electric field in the target region has been common (Axel Thielscher et al., 2015). Our findings imply that such simulations may also be beneficial in predicting the effectiveness of tSMS. An important avenue for future research with tSMS might be to control the dosage across participants based on these simulations, allowing for an optimal selection of magnetic strength that standardizes the dose or effectiveness of tSMS across different participants and target regions.

In the present study, the baseline block was conducted first, followed by the test block with the magnet in place. One might argue that this fixed order could introduce a potential confound, as task performance may improve over time due to learning effects. However, several lines of evidence suggest that the improvement observed in our study cannot be fully explained by learning alone. First, if practice were the primary driver of the observed improvement, similar effects would be expected regardless of the stimulation site; however, no such improvement was observed in the SMA condition. Second, by subtracting baseline thresholds within each session, the influence of learning across sessions was minimized. Finally, individual differences in threshold improvement observed in the IPL condition were correlated with scalp-to-IPL distance, which cannot be accounted for by simple practice effects and instead supports the presence of a physiological impact of tSMS. Taken together, these findings strengthen the interpretation that the observed enhancement reflects a site-specific physiological effect of tSMS rather than a learning-related effect.

### Limitations

A limitation of the current study is that our experimental design, which employed the staircase method, only assessed the modulation of threshold. Consequently, it remains unclear whether tSMS impacted perceived duration and/or orientation (i.e., bias). Previous studies have suggested that the right IPL is associated with the subjective experience of time (Wiener et al., 2010a, 2012; Hayashi and Ivry, 2020), highlighting the necessity of measuring biases in perceived duration. This can be clarified by using discrimination tasks that involve a stable reference unaffected by tSMS, such as a bisection task with an internal reference.

Second, since tSMS was applied continuously during the discrimination tasks, the specific processes being modulated (e.g., stimulus encoding, maintenance, comparison) in the orientation task remain unclear. This lack of temporal specificity may not be easily addressed due to the nature of tSMS, necessitating the use of more temporally specific stimulation methods, such as focused ultrasound stimulation.

Finally, the absence of a sham condition represents an additional limitation of the present study. The inclusion of a sham magnet condition would allow for more rigorous control of nonspecific influences, including practice-related and placebo effects. Future studies incorporating such a sham condition would help fully dissociate the specific physiological effects of tSMS from these potential confounds.

### Conclusions

The current study aimed to identify the brain regions that are causally involved in temporal and spatial perception using tSMS. The observed improvements in orientation discrimination thresholds induced by tSMS suggest a potential causal involvement of the right IPL in spatial processing. Furthermore, we demonstrated that individual differences in the distance from the scalp to the right IPL correlated with the effectiveness of tSMS, underscoring the importance of considering anatomical differences when predicting and evaluating the efficacy of tSMS.

## CRediT authorship contribution statement

**Masakazu Sugimoto:** Methodology, Software, Validation, Formal analysis, Investigation, Data Curation, Writing-Original draft preparation, Visualization. **Ikko Kimura:** Methodology, Writing-Reviewing and Editing. **Masamichi J. Hayashi:** Conceptualization, Methodology, Software, Writing-Reviewing and Editing, Supervision, Project administration, Funding acquisition

## Acknowledgments

This work was supported by the Japan Society for the Promotion of Science (Grants-in-Aid for Scientific Research JP21H00315, JP22H01110, and JP23K17649 to M.J.H., and Grant-in-Aid for JSPS Fellows JP25KJ1802 to M.S.) and the Japan Science and Technology Agency (FOREST JPMJFR232X to M.J.H.).

## Declaration of generative AI and AI-assisted technologies in the writing process

During the preparation of this work, the authors used ChatGPT, Grammarly, and DeepL in order to improve the readability and language of the manuscript. After using these tools, the authors reviewed and edited the content as needed and took full responsibility for the content of the published article.

